# Model communities hint to promiscuous metabolic linkages between ubiquitous free-living freshwater bacteria

**DOI:** 10.1101/103838

**Authors:** Sarahi L Garcia, Moritz Buck, Joshua J. Hamilton, Christian Wurzbacher, Magnus Alm Rosenblad, Katherine D. McMahon, Hans-Peter Grossart, Falk Warnecke, Alexander Eiler

## Abstract

Free-living microorganisms with streamlined genomes are very abundant in the environment. Genome streamlining results in losses in the cell’s biosynthetic potential generating physiological dependencies between microorganisms. However, there exists no consensus on the specificity of these microbial associations. To verify specificity and extent of these associations, mixed cultures were established from three different freshwater environments. These cultures contained free-living streamlined organisms lacking multiple biosynthetic pathways. Among the co-occurring members of the mixed cultures, there was no clear recurring pattern of metabolic complementarity and dependencies. This, together with weak temporal co-occurrence patterns observed using time-series metagenomics, suggests that free-living freshwater bacteria form loose and unspecific cooperative loops. Comparative genomics suggests that the proportion of accessory genes in populations of streamlined bacteria allows for flexibility in interaction partners. Altogether this renders these free-living bacterial lineages functionally versatile despite their streamlining tendencies.

## Introduction

Microorganisms can cooperate in many different ways and their relationships range from facultative to obligate dependencies ^1^. On the far end of the dependency spectrum are endosymbionts (and endoparasites). These show obligate dependencies where genome reduction and associated loss of essential biosynthesis pathways are widespread. In most cases, a defined host - endosymbiont specificity is established. At the other end of the spectrum, free-living bacteria are considered to be widely autonomous. However, this image is now changing rapidly with frequent reports of reduced microbial genomes in the environment resulting in unique and singular auxotrophies ^2^.

In natural aquatic environments where nutrients generally occur in low concentrations, microbes produce many compounds that are costly, but promote survival and reproduction not only for themselves but also for neighboring cells ^3^. In fact, auxotrophy for amino acids and vitamins has recently been shown for numerous free-living bacteria ^4–6^, pointing to critically important metabolic dependencies from other community members as exemplified in a simple model community ^7^. Therefore, it would not be surprising that most natural aquatic systems allow for frequent and complex metabolic interactions, for example, via continuous mixing of the environment and molecular-scale diffusion facilitating distribution of public goods. Hence, even when abundant environmental bacteria are seen as free-living, their streamlined genomes render them tightly linked and dependent on other microorganisms ^8^ in the community. The sheer number of bacteria and hence genetic variability in a population could allow for high metabolic flexibility at the population level with an endless number of metabolic interactions possible among free-living bacterial communities. However, the character of these associations still remains largely unknown.

To investigate the specificity of the dependencies between free-living freshwater microorganisms, we established dilution mixed cultures as model communities. These cultures are a small subsample of the complex natural community ^9^ with about 12 initial cells. About 600 of such mixed cultures were established and screened for a group of cosmopolitan yet streamlined freshwater organisms from the Actinobacterial lineage acI. Here, we describe six of these model communities and resolve their metabolic interaction potentials using metagenomics. Finally, for a single mixed culture, we calculate co-occurrence patterns across a metagenomics time-series in a lake to test if the observed dependencies are specific or if the interaction-partners are flexible in more complex natural communities.

## Results and Discussion

### Mixed cultures as model communities and their genomic reconstructions

Ubiquitous freshwater Actinobacteria of the acI lineage were cultivated by diluting different lake samples with triple-filtered sample water incubated in 96-well liquid culture plates. In total, seven culture plates were set up and around forty-five wells contained acI. Of these, only 6 cultures successfully propagated with densities comparable to those observed in the environment (i.e. 10^6^ cells ml^-1^) for subsequent transfers and growth in larger volumes. This resulted in 6 dilution-to-extinction mixed cultures: FNEF8, FNEB6, FNEB7, FNED7, FSWF8 and TBE6 (FNE refers to Northeast basin of Lake Grosse Fuchskuhle, FSW to Southeast basin of Lake Grosse Fuchskuhle and TB to Trout Bog Lake). Characterizations of FNEF8 have been published, previously ^7,10^.

DNA was extracted from 4 L-cultures, and reads obtained by shotgun sequencing were assembled. Contigs > 1000 bp were considered for further analysis. In total, 77 metagenome assembled genomes (MAGs) were obtained from the assemblies (Supplemental). Of these, 31 MAGs each recruited more than 1% of the reads in the original metagenome. These MAGs have an average completeness of 83.5% as calculated by CheckM (Supplemental). It has been previously observed that because CheckM relies on lineage-specific marker genes, the completeness of genomes without lineage representation can often be underestimated ^5,7^. However, since these MAGs have extremely high coverage and were assembled from model communities (low diversity of starter cells) we are confident of their high quality.

### Some free-living microorganisms fulfill their metabolic needs from those they happen to encounter

Populations represented by the top 31 MAGs were assumed to be the dominant model community members and were analyzed further (mapping rates in Supplemental). All six cultures yielded at least one MAG of the cosmopolitan freshwater Actinobacteria acI (recruiting between 15% and 40% of reads), four cultures yielded a MAG of the freshwater Actinobacteria acIII (between 3% and 9% of the reads), two cultures yielded yeast MAGs (1 and 3% of the reads), two cultures featured MAGs affiliated with Bacteroidetes (2% and 6% of the reads), two cultures yielded a Polynucleobacter MAG (3% and 55% of the reads), two cultures yielded a Spirochaetes MAG (8% and 22% of the reads), one culture yielded an Alphaproteobacteria MAG (1% of the reads) and one an Acidimicrobiales MAG (48% of the reads). Knowing that acI Actinobacteria do not grow in pure culture ^11^, we now have strong support that the free-living acI is highly dependent on interactions with other microorganisms. Results from previous co-occurrence studies support the idea that free-living streamlined bacteria have very high connectivity in their environments and are critically dependent on metabolites that might be provided by the surrounding members of the planktonic communities ^8,12^. Moreover, by creating an initial bottleneck ^13^ of approximately 12 cells, we obtained a first evidence that these plankton community partners do represent a diverse sets of community members. This hints towards a non-specific metabolic dependence of acI on other abundant freshwater bacteria, as no consistent interaction patterns were observed. Since six mixed cultures might be not sufficient to generalize the character of the observed associations, we correlated the abundance of the mixed culture TBE6 MAGs with those obtained from a nine-year time-series shotgun metagenome from its source environment, Trout Bog Lake ^14^. For this, we also used MAGs reassembled from this time-series data. The observed correlations indicate that acI interacts promiscuously and randomly (Figure 1, Figure S1 and S2). That is, no strong exclusive correlations can be discerned, either with the members of TBE6 or MAGs from the time-series, strengthening the hypothesis that acI is promiscuous in its interactions.

**Figure 1.**
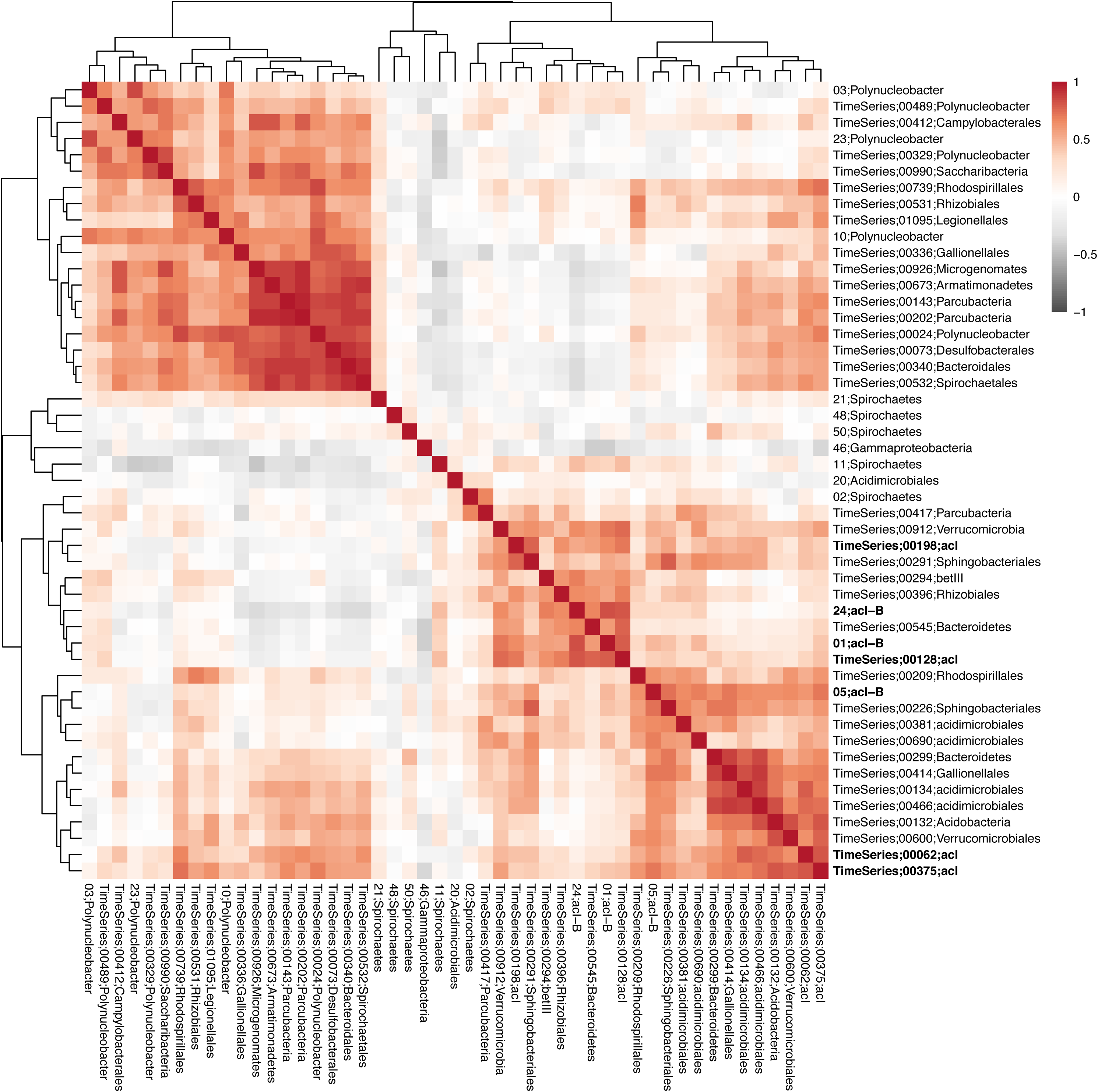
Spearman correlation on normalized relative abundance between MAGs from TBE6 and taxa in the epilimnion of the environment of origin (Trout Bog Lake). Metagenome samples from time-points from all years were used (45 unique dates). Correlation is displayed via hierarchical clustering. MAG names that start with a number refers to a MAG from the mixed culture (for details on these MAGs look at supplemental tables). MAG names that start with the words “time series” were previously binned from the Trout Bog nine year metagenome series ^14^. The acI MAGs are highlighted in bold. To look into the correlations of hypolimnion and both combined look into the supplemental.

### Tight metabolic dependencies between different free-living microorganisms

Canonical central metabolic pathways such as glycolysis, the pentose phosphate pathway, the citric acid cycle, ammonia assimilation and oxidative phosphorylation are observed in each of the abundant MAGs of the microbial model communities, suggesting a mainly aerobic heterotrophic lifestyle (Supplemental). When looking at all extant acI MAGs and single-cell genomes (SAGs) ^15,16^, auxotrophies for amino acids and vitamins were observed (Figure 2A). However, in the mixed cultures, at least one of the main members in each community was able to synthesize each of these metabolites, hinting to a high degree of metabolic dependency within the members of each model community of relatively low complexity. For example, in culture FNEB6, Bacteroidetes is the only producer of asparagine and so potentially supplies it to the other community members (see more examples in Figure 2B). Interactions at the level of amino acids and vitamins have been previously observed and described in several aquatic study systems ^4–6^. Furthermore, one or two members of each model community had the ability to reduce sulfate, consistent with prior findings of the transfer of redox reaction products known as “metabolic handoffs” ^17^. These kinds of observed dependencies merely represent a few examples of the many interactions that likely occur in the complex natural environment. In this respect, it was interesting that they also formed cross-domain linkages to two ubiquitous yeasts. We consider this as an indication that these interactions potentially can cross boundaries to eukaryotic plankton.

**Figure 2.**
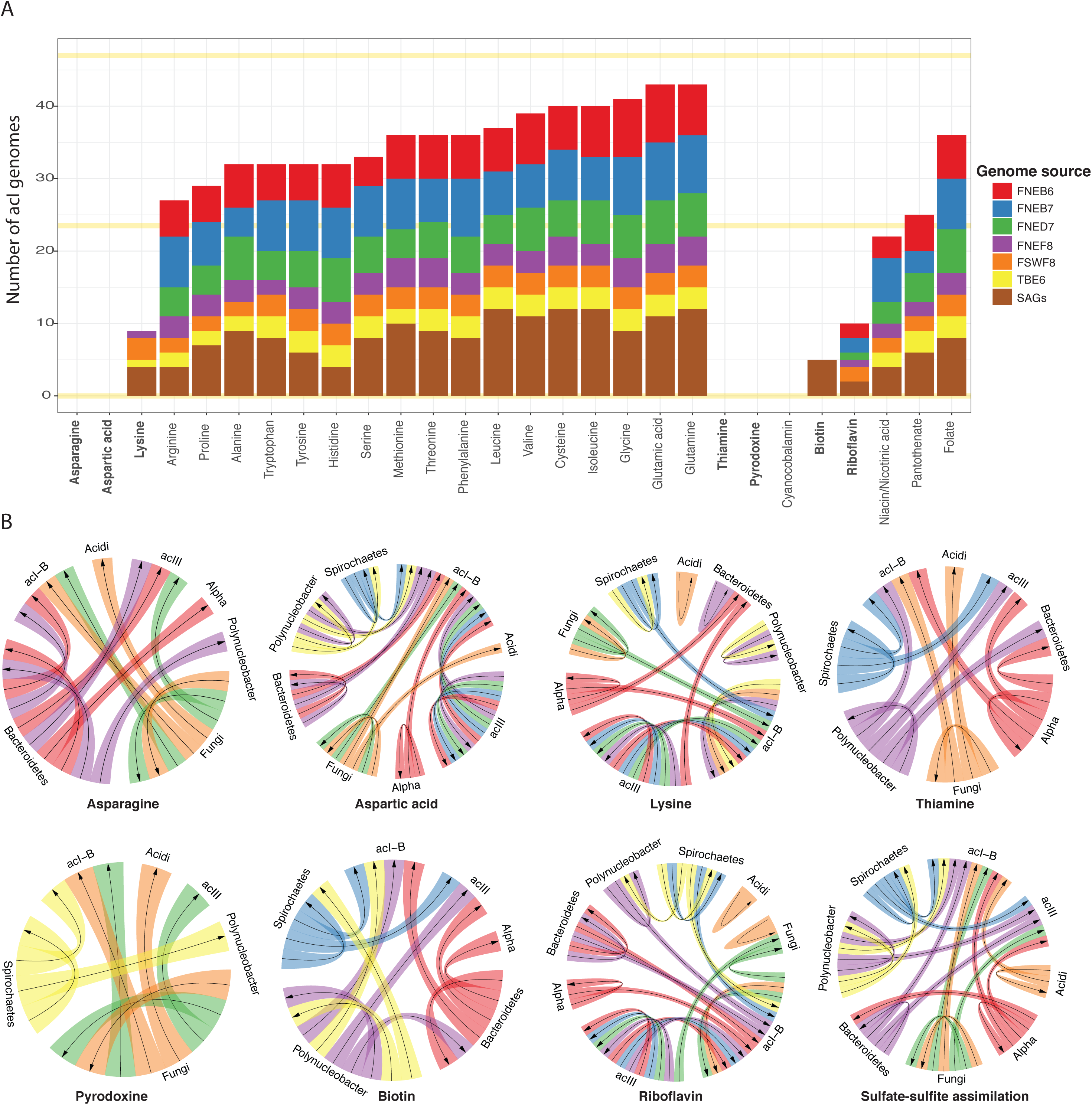
**A.** Visualization of the amino acid and vitamin biosynthetic potential of all acI MAGs and SAGs analyzed. It includes acI information from all MAGs binned from each culture in this study plus the previously published SAGs. Yellow lines indicate 0%, 50% and 100% of genomes analyzed. Completeness of a metabolic pathway was considered if 80% of the pathway was present. **B.** Potential metabolic complementarity among major members of each of the cultures. Circle plots are displayed for metabolites highlighted in bold in panel A plus potential to assimilate sulfate and sulfite. Colors indicate each mixed culture using same color scheme as in panel A. Arrow shows the direction in which the metabolite would be “shared”. Alpha – Alphaproteobacterium. Acidi – Acidimicrobium.

### High degree of genomic diversity might support flexibility in interaction partners

Our findings from analysis of genome content combined with prior evidence of high genome-level diversity within acI ^15^ prompted us to examine the variability in gene content among acI genomes. Interestingly, even when our cultures harbor a reduced diversity, all of our cultures contained more than one genotype of acI. This likely reflects the large genomic plasticity of this common and dominant free-living freshwater bacterium. This genomic plasticity, however, cannot be resolved with the short 16S rRNA gene fragments obtained in normal amplicon sequencing (Supplemental). Using all 33 acI Actinobacteria MAGs from our 6 mixed cultures in addition to the 14 available acI single cell amplified genomes (SAGs) (Figure S3), ~8000 orthologous gene clusters (OGCs) were found. The distribution of gene clusters in the genomes revealed that ~800 OGCs form the core genome of acI (Figure S4 and S5). Assuming around 1600 genes per acI ^
15^, this means that about half of the whole acI genome belongs to the flexible genome. This is around the same proportion as SAR11 ^
18^. Thus, both acI and SAR11 have quite larger flexible genomes than other ubiquitous free-living aquatic bacteria like Prochlorococcus ^19^. The paradox of acI's streamlining tendencies is that their high number of auxiliary genes is likely to render populations of this free-living bacterial lineage functionally versatile.

In summary, as is the case for other streamlined free-living aquatic bacteria, acI Actinobacteria depend on numerous other abundant microorganisms for metabolic handouts. This kind of dependence seems to be non-taxa-specific or promiscuous since no consistent interaction patterns could be observed. These observations add to the knowledge on the structure of cooperation and dependencies between free-living aquatic microbes.

## Methods

### Mixed culture and DNA extraction

Mixed cultures that included the abundant and ubiquitous freshwater *Actinobacteria* were established in 2012 and 2013. Samples were collected from 5 different lakes: Lake Grosse Fuchskuhle (North East basin and South West basin), Lake Stechlin, Lake Dagow, Lake Mendota and from Trout Bog Lake (metadata reported in Supplementary Table). In total, seven plates were set up, two for Northeast basin (Lake Grosse Fuchskuhle) and one for each of the other lakes (Southwest basin Lake Grosse, Lake Stechlin Lake, Lake Dagow, Lake Mendota and Trout Bog Lake. Cultivation method, media preparation, acI qPCR assays, general observations and DNA extractions were as previously described for mixed culture FNE-F8 ^10^. Selected mixed cultures (based on cell density) were scaled up to a four-liter culture and DNA was extracted. DNA was sent to JGI and to Research and Testing Laboratory (RTL) (http://researchandtesting.com/) for shotgun metagenomic sequencing and 16S rRNA gene amplicon sequencing, respectively.

### 16S rRNA gene amplicon sequencing

RTL performed PCR amplification and MiSeq sequencing of the 16S rRNA gene amplicons. The bacterial primers (*E. coli* positions) were 357wF (5’-CCTACGGGNGGCWGCAG-3’) and 926wR (5’-CCGTCAATTYMTTTRAGTTT-3’). All downstream analysis was done using the RTL standard pipeline from 2014. In brief all sequences were clustered at 4% divergence into OTUs using the UPARSE ^20^ algorithm. The centroid sequence from each cluster was then run against either the USEARCH global alignment algorithm or the RDP Classifier against a database of high quality sequences derived from the NCBI database. The output was then analyzed using an RTL internally developed python program that assigns taxonomic information to each sequence and then computes and writes the final analysis files.

### Library preparation and sequencing

JGI performed the library preparation and the sequencing under Community Sequencing Project 1289. First, 100 ng of genomic DNA was sheared to 270 bp using a focused-ultrasonicator (Covaris). The sheared DNA fragments were size selected using SPRI beads (Beckman Coulter). The selected fragments were then end-repaired, A-tailed, and ligated of Illumina compatible adapters (IDT, Inc.) using KAPA-Illumina library creation kit (KAPA biosystems). The prepared sample library was quantified using KAPA Biosystem’s next-generation sequencing library qPCR kit and run on a Roche LightCycler 480 real-time PCR instrument. The quantified library was then prepared for sequencing on the Illumina HiSeq sequencing platform utilizing a TruSeq paired-end cluster kit, v3, and Illumina’s cBot instrument to generate a clustered flowcell for sequencing. Sequencing of the flowcell was performed on the Illumina HiSeq2000 sequencer using Illumina TruSeq SBS sequencing kits, v3, following a 2x150 indexed high-output run recipe.

### Genome assembly, annotations and metabolic features analysis

After reads had been filtered based on their quality scores using sickle (version 1.210) ^21^, the reads were digitally normalized using khmer 1.4 ^22^ and assembled with megahit ^23^. Coverage was computed by mapping back the reads competitively to the assembly using bbmap 35.40 (sourceforge.net/projects/bbmap/) using default parameters. Mapped reads were indexed and sorted using SAMtools 1.3 ^
24^ while we removed duplicates using picard-tools (version 1.101). Bedtools (version 2.18.2) ^25
^ was used for computing coverage. For the binning metaBAT ^26^ was used. Tribe/taxonomical affiliation were defined using whole genome information and the public database using PhyloPhlAn ^27^ and using previously published genomes and SAGs ^16^ as references. If the MAGs belong to one of the freshwater tribes ^28^ they have the name of the tribe, otherwise the taxonomic name after NCBI is given.

For bacterial genomes, gene prediction analysis was performed within the Integrated Microbial Genomes (IMG) ^29,30^ platform developed by the Joint Genome Institute, Walnut Creek, CA, USA (http://img.jgi.doe.gov). Specific KEGG biosynthetic maps were inspected for completeness (http://www.genome.jp/kegg/mapper.html) counting number of missing enzymes from the most complete pathway. MAG completeness was calculated using CheckM ^31^.

For fungal genomes, ab initio gene prediction was performed using augustus (v 2.5.5) with Aspergillus nidulans (FSWF8-4) or Cryptococcus neoformans (FNED7-22) models. Gene model ORFs were then annotated using matches to Pfam 30.0 (June 2016) by hmmscan (HMMER 3.1b2). Functional annotation was done using predicted aminoacids in the KAAS platform ^32^.

### Core genome computation

Clusters of orthologous genes were identified using OrthoMCL ^33^ using default options following an established protocol. Core genome of the acI was computed using all 33 acI Actinobacteria MAGs from our 6 mixed cultures in addition to the 14 available acI single cell amplified genomes (SAGs). Due to the incompleteness SAGs and MAGs, common methods, such as picking all the OGCs present in at least 90% of the genomes, are unsafe. To reduce the noise generated by incompleteness, a semi-supervised approach has been chosen. For this we compute a distance matrix based on the genomes that contain the different OGCs. Two OGCs will have a distance of 0.0 if they are found in the same genomes, and a distance of 1.0 if none of the genomes containing one also contains the other. The overlap between the genome sets of each cluster were computed and normalized to the smallest genome set. OGCs contained in less than 4 genomes were removed. The obtained distances are then used with a hierarchical clustering to compute a heatmap (Supplemental Figure acI core). This heatmap clearly shows a set of OGCs that overlaps significantly with all other OGCs, this cluster is selected as core. Core OGCs should have a strong overlap with all other OGCs irrelevant of their position in the heatmap.

### Metagenome recruitment to MAGs

A total of 113 metagenome samples from Trout Bog lake ^14^ were used in this study. As above, after assembly and binning, bbmap was used to compute the coverage of all the MAGs (from the time-series as well as the cultures), for all the time-points. All coverages have been normalized to sequencing depth. Mean coverage vectors of all MAGs are then correlated using Spearman correlation. For the heatmaps, MAGs from Trout Bog lake were only kept if they correlate with at least 0.6 to at least one MAG from our mixed culture TBE6 (positive or negative correlation). MAGs assembled from Trout Bog lake have not been further analyzed as they might represent an average population genome.

## Acknowledgements

In memory of Falk Warnecke, who believed in the power of mixed cultures and envisioned this manuscript already a few years ago. This work was supported by: the Swedish research council VR (grant 2012-4592), the German Science Foundation (grants DFG GR1540/ 17-1 and 21-1), the JSMC, the United States National Science Foundation Microbial Observatories program (MCB-0702395), the Long Term Ecological Research program (NTL-LTER DEB-1440297), and INSPIRE award (DEB- 1344254). JJH was supported by a USDA NIFA AFRI Postdoctoral Fellowship, award number 2016-67012-24709. We are also grateful to the Joint Genome Institute for supporting this work through the Community Sequencing Program (CSP) and providing technical support. The work conducted by the U.S. Department of Energy Joint Genome Institute, a DOE Office of Science User Facility, is supported by the Office of Science of the U.S. Department of Energy under Contract No. DE-AC02-05CH11231. Thanks to Alexandra Linz for her support with the TBE6 mixed culture. Thanks to staff at the Joint Genome Institute including Rex Malmstrom, Matthew Bendall, Tijana Glavina del Rio, Rob Eagan, and Susannah Tringe.

## Data accessibility

The raw shotgun metagenome reads are publicly available in the JGI portal and the assembly is available in IMG/MER under the ER submission IDs 26656, 26658, 26650, 29729, 29808, 50227. The bacterial metagenome-assembled genomes (MAGs) are also available through IMG. The fungal MAGs have been deposited at DDBJ/ENA/GenBank. For taxon OID or accession number check supplementary material.

## Author contribution

SLG and FW conceived the research. SLG and HPG collected and prepared the samples. SLG, MB, JJH, AE, CW and MAR analyzed the data. All authors wrote and/or revised the manuscript.

